# Action features dominate cortical representation during natural vision

**DOI:** 10.1101/2025.01.30.635800

**Authors:** Heejung Jung, Xiaochun Han, Ma Feilong, Jane Han, Deepanshi Shokeen, Cara Van Uden, Isabella Hanson, Andrew C. Connolly, James V. Haxby, Samuel A. Nastase

## Abstract

Cortical resources are allocated into systems specialized for processing ecologically relevant features of the world. These features are typically studied in isolation using controlled experimental stimuli, making it difficult to assess the relative importance of different kinds of features that tend to overlap during natural vision. In the current study, we evaluated the relative contributions of action, agent, and scene features in predicting cortical activity while participants viewed a 1-hour nature documentary. We tested four sets of model features: semantic vectors capturing the observed action, agent, and scene features derived from an annotation of the stimulus, as well as low-level visual motion-energy features. We used banded ridge regression to fit vertex-wise encoding models from the combination of all four sets of features. While each set of features predicted neural activity in the expected areas, the action features predicted more widespread activity. A variance partitioning analysis revealed that the action features captured the most unique variance in neural activity across ten times as many cortical vertices as the agent or scene features. Our findings suggest that cortical activity during dynamic, natural vision is dominated by features for understanding the actions of others.

## Introduction

Cortical resources are differently allocated toward processing different features of the world. This division of labor coincides with the behavioral relevance of certain kinds of stimuli. For example, in highly visual species like humans and other primates, a relatively large proportion of cortex is devoted to visual processing, and visual cortex is highly subdivided, relative to other mammals (Kaas, 1989, 2004; Felleman & Van Essen, 1991). Cortical magnification as a function of behavioral importance and complexity is a well-established principle in sensory and motor cortices: foveal vision recruits much more cortex than the periphery for higher-acuity processing (Daniel & Whitteridge, 1961; Cowey & Rolls, 1974; Rovamo et al., 1978; Malach et al., 2002; Duncan & Boynton, 2003); effectors requiring fine motor control like the hands and mouth occupy disproportionately large portions of sensorimotor cortex (Penfield & Boldrey, 1937; Woolsey et al., 1942; Penfield & Rasmussen, 1950; Nudo et al., 1992; Pascual-Leone & Torres, 1993). Cortical magnification may also extend into higher-order perceptual and association cortices, reflecting the relative complexity and ecological importance of different high-level features.

In this vein, functional neuroimaging has revealed a mosaic of functionally specialized cortical areas for processing faces (Kanwisher et al., 1997), scenes (Epstein & Kanwisher, 1998), and other salient features of our visual world (Kanwisher, 2010), culminating in a ventral visual pathway for object perception and categorization (Kravitz et al., 2013; Grill-Spector & Weiner, 2014). Results of this kind have informed a number of theories on the large-scale structure of semantic knowledge, particularly object categories, in the human brain (Barsalou et al., 2003; Caramazza & Mahon, 2003; Martin, 2007, 2016; Patterson et al., 2007; Mahon & Caramazza, 2009). In these experiments, however, neural responses to static images of faces or scenes are typically contrasted with carefully selected control images matched on certain parameters—making it difficult to assess how different kinds of features compare in terms of their overall cortical footprint.

A largely separate line of work has focused on the cortical systems supporting the perception of dynamic visual features, such as biological motion (Bonda et al., 1996; Grossman et al., 2000; Beauchamp et al., 2003; Puce & Perrett, 2003; Gobbini et al., 2007) and observed actions (Castelli et al., 2000; Gobbini et al., 2007; Grafton & Hamilton, 2007; Oosterhof et al., 2013; Lingnau & Downing, 2015). These experiments typically employ highly-controlled stimuli as well: for example, point-light displays of a figure moving naturally versus non-natural motion (e.g., Grossman & Blake, 2002; Saygin et al., 2004), moving geometric shapes depicting social interactions (e.g., Castelli et al., 2000; Gobbini et al., 2007), or brief videos depicting isolated hands performing simple actions like opening versus closing a container (e.g., Oosterhof et al., 2012; Wurm & Lingnau, 2015). Experiments of this kind have culminated in a lateral visual pathway for dynamic social perception (Pitcher & Ungerleider, 2021; McMahon & Isik, 2023) and a frontoparietal action observation network (Grafton & Hamilton, 2007; Rizzolatti & Sinigaglia, 2010; Oosterhof et al., 2013). Ventral visual areas, however, are also consistently activated by these dynamic stimuli depicting agentic action (e.g., Bonda et al., 1996; Castelli et al., 2000; Gobbini et al., 2007), suggesting that the putative “object vision pathway” also plays an important role in dynamic social perception and action observation (Haxby, Gobbini, et al., 2020; Han et al., 2024). In studies where different classes of stimuli are compared (e.g., Haxby et al., 2001; Beauchamp et al., 2002; Grossman & Blake, 2002), many regions exhibit mixed responses to different classes of stimuli, suggesting that functional specialization may be more complex than some experiments would suggest.

These cortical pathways for static object perception and dynamic action perception emerged separately, in isolated experiments. Taking these findings into consideration, we might surmise that, while cortex has functionally specialized subdomains for certain features of the world, these features are roughly commensurate—each of these features is similarly important to cortex. This conclusion may not be warranted, however, for several reasons. While targeted experimental manipulations are useful for disentangling different features, this piecemeal approach limits our ability to quantify how different features are processed in cortex under naturalistic conditions, when the field is cluttered, dynamic, and embedded in a temporal and semantic context. In natural vision, features are not compartmentalized—they are often superimposed and may interact in a number of ways. For instance, different agents may perform semantically similar actions in kinematically different ways (a monkey eating nuts, a seal eating penguins); different actions may correlate with certain background features (e.g., a cheetah sprinting across the savanna, a dolphin swimming past a reef). While these might be considered confounds in experimental design, the brain may both capitalize on these statistical associations (Olshausen & Field, 1996; Simoncelli & Olshausen, 2001; David et al., 2004; Theunissen & Elie, 2014; Hasson et al., 2020) and learn invariances across them: despite the fact that a caterpillar, a hummingbird, and a chimpanzee *eat* in quite different ways, the brain encodes semantic features of the *eat* action that generalize across different agents (Nastase et al., 2017). The use of naturalistic stimuli allows us to more holistically evaluate the relative contribution of different features that are typically studied in isolation (Hamilton & Huth, 2020; Haxby et al., 2020; Nastase et al., 2020).

Recent work has begun using movies and spoken narratives to explore how different features are encoded in neural activity under more ecological conditions. Movies have been shown to drive reliable activity across much of cortex (Bartels & Zeki, 2004; Hasson et al., 2004, 2010; Jääskeläinen et al., 2008; Haxby et al., 2011; Baldassano et al., 2017; Kim et al., 2018; Richardson et al., 2018; Yang et al., 2023; Rajimehr et al., 2024) and are thought to sample features of the world (and their intercorrelations) in a more “representative” (Brunswik, 1955) way than typical experimental stimuli (Sonkusare et al., 2019; Nastase et al., 2020). For example, Huth and colleagues (2012) used an annotation of a natural movie stimuli to demonstrate that different semantic features are intermixed and represented broadly across cortex. Unexpected results have been found in macaques, where dynamic, socially-relevant features of naturalistic stimuli were found to drive activity in areas typically associated with face processing in experiments using static image stimuli (Russ & Leopold, 2015; Russ et al., 2023). Given the complexity inherent to naturalistic stimuli, how can we assess what features best capture cortical activity? Researchers have developed a modeling framework where the stimulus is decomposed into different feature spaces based on explicit models, which are then tested against the neural activity using joint and nested regression models (Lescroart et al., 2015; de Heer et al., 2017; Lee Masson & Isik, 2021). For example, de Heer and colleagues (2017) compared three different sets of features for speech comprehension in spoken narratives: spectral, articulatory, and semantic features. Using a variance partitioning analysis, they found that semantic features capture much more variance in neural activity across a large portion of cortex, whereas spectral and articulatory features of speech were encoded in relatively focal auditory regions. This framework allows us to quantitatively link complex neural activity driven by naturalistic stimuli to interpretable features, and compare the relative contributions of different types of features.

In the current study, we asked whether certain higher-level features of the visual world are “magnified” across cortex relative to other seemingly commensurate features. We tested several different kinds of ecologically-relevant visual features to determine whether any particular set of features recruits a disproportionately large amount of cortical resources. We measured brain activity using fMRI while subjects viewed a 1-hour movie—the *Life* nature documentary narrated by David Attenborough—depicting different animals behaving in their natural environments. Based on predominant trends in cortical organization emerging from the literature, we tested three different sets of features capturing the semantic content of the *actions* (e.g., swimming, running), *agents* (e.g., dolphin, cheetah), and *scenes* (e.g., ocean, savanna) visually depicted over the course of the film. We also include a motion-energy model to capture the low-level visual motion of the stimulus and better isolate higher-level semantic features. To compare these features, we estimated vertex-wise encoding models jointly across all four feature spaces, then assessed the relative contribution of each feature space. We found that the action features predominate in dorsal regions, with strong encoding performance extending from inferior parietal regions to premotor cortex. The agent features yielded the strongest encoding performance in ventrolateral temporal cortex, extending from lateral fusiform to the posterior lateral occipitotemporal cortex. The scene features were strongest in ventromedial occipital and temporal cortex. A variance partitioning analysis, however, revealed that the action features uniquely predicted activity across a much greater proportion of cortex than the competing models.

## Results

We tested the relative contributions of different semantic feature spaces to cortical activity while participants viewed a nature documentary (Fig. 1). Semantic features for *actions*, *agents*, and *scenes* were constructed from a detailed behavioral annotation of the visual content of the film (Huth et al., 2012). For each word, we obtained a 300-dimensional word embedding from the GloVe model (Pennington et al., 2014). In this model, semantic relationships are encoded in the geometric relationships between vectors, such that more similar words are located nearer to each other in the embedding space, based on their co-occurrence in large corpora of text. We also tested a biologically-inspired motion-energy model capturing dynamic, low-level visual content (Adelson & Bergen, 1985; Watson & Ahumada, 1985; Nishimoto et al., 2011). To assess these different models, we constructed vertex-wise encoding models: we fit a linear model mapping from the semantic vectors to the response time series at each vertex (Naselaris et al., 2011). To fairly assess the contribution of each model, we first used principal component analysis (PCA) to reduce each feature space to a matching dimensionality (40 dimensions, explaining over 95% of the variance in each of the three feature sets of interest; Fig. S1). Second, we combined all four feature spaces and estimated joint encoding models using banded ridge regression (Dupré la Tour et al., 2022). We evaluated the relative contributions of these models using four-fold, leave-one-run-out cross-validation combined with leave-one-subject-out cross-validation; that is, models were evaluated in a left-out segment of the stimulus in a left-out subject. To better align cortical-functional topographies across individuals, we estimated individual-specific transformations using whole-brain surface-based searchlight hyperalignment (Guntupalli et al., 2016) from the three training runs, then applied these transformations to the left-out fourth run. We quantified encoding performance as the correlation between the model-predicted time series and the actual fMRI time series for the test run in the test subject.

**Figure 1.**
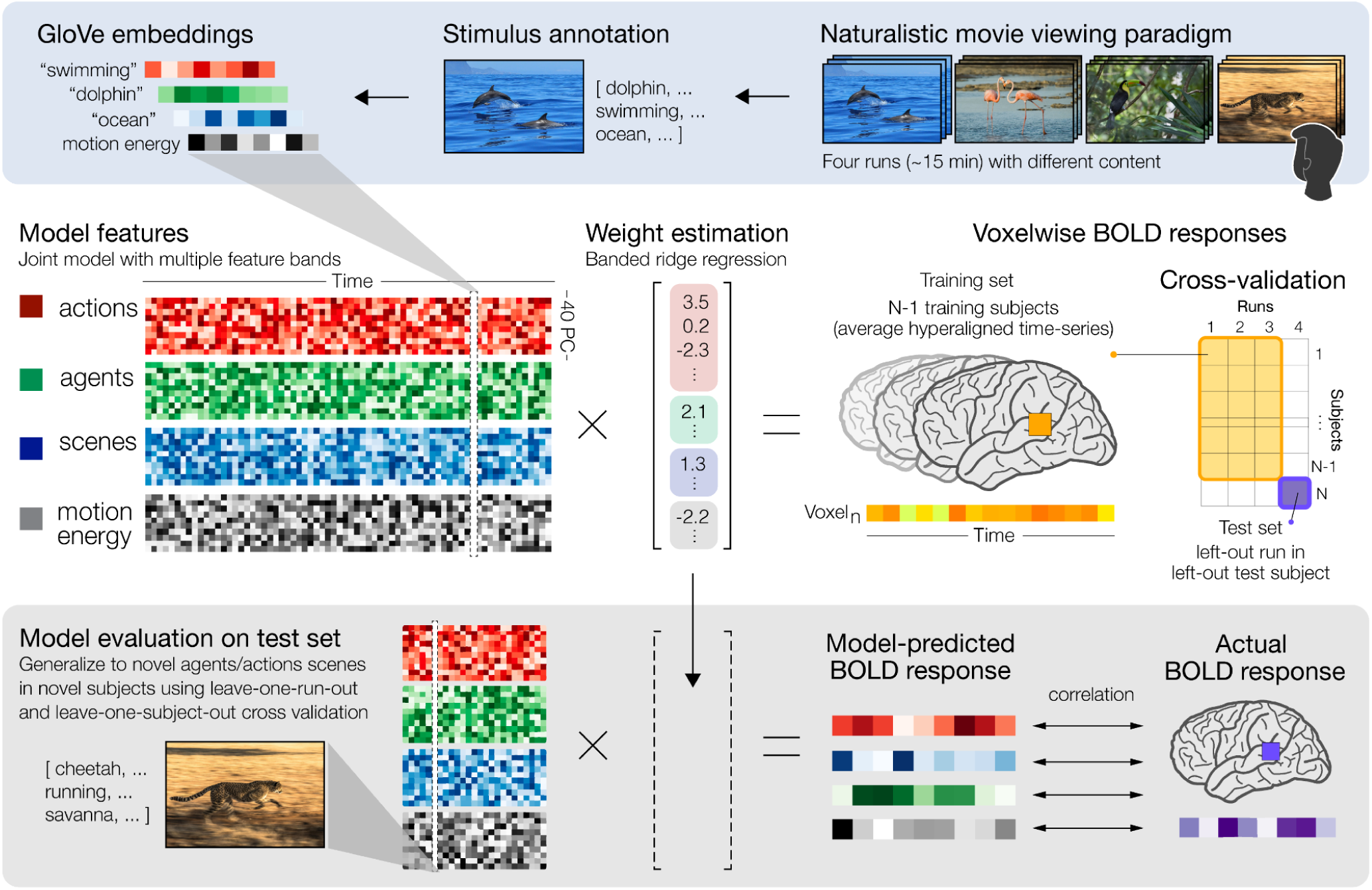
Encoding multiple feature spaces using banded ridge regression. Participants watched a movie stimulus—a nature documentary featuring a variety of behaving animals in their natural environments—during fMRI acquisition. The stimulus was split into four separate ∼15-minute runs; each run was unique and contained multiple scenes depicting different animals in different environments. The stimulus was annotated at roughly 2-TR (5-second) intervals to describe in words what was visually depicted in the film. The annotation was then split into words describing the agents (e.g., dolphin), actions (e.g., swimming), and scene (e.g., ocean). For each word, we extracted GloVe embeddings capturing the semantic content of the words. We also extracted dynamic, low-level visual features of the stimulus using a motion-energy model. The embeddings for agent, action, and scene words, as well as the frame-by-frame motion energy features, were downsampled to the TR and concatenated into a joint encoding model. We used banded ridge regression to estimate a linear mapping from the joint model to the BOLD response time series at each vertex. Vertex-wise encoding models were trained on the average time series from *N* – 1 participants aligned into a shared space using searchlight hyperalignment (leave-one-subject-out cross-validation) for three of the four stimulus segments (leave-one-run-out cross-validation). We used the trained model to generate predicted BOLD response time series for the test segment. Model-based predictions were generated jointly or separately for each feature space by discarding (zeroing out) weights for other feature spaces. Encoding model performance was evaluated by computing the correlation between the model-predicted BOLD time series and the actual BOLD time series for the left-out test run in the left-out test subject (also aligned to the shared space using hyperalignment transformations based on the training runs).

### Vertex-wise encoding performance for actions, agents, and scene features

To quantify the relative contributions of each feature space, we fit joint vertex-wise encoding models, then generated four sets of predictions based on the subset of weights corresponding to each feature space. We computed the correlation between the predictions for each feature space and the actual vertex time series separately (see Fig. S2 for the full model performance). All four feature spaces yielded significant predictions (*t*-test, FDR *p* < .05) with different cortical topographies (Fig. 2). The action features yielded significant predictions throughout the action observation network extending from lateral occipitotemporal cortex (LOC), anterior intraparietal (AIP) and postcentral cortex, and premotor cortex (Fig. 2a). Predictions based on the agent features were strongest in LOC, posterior superior temporal cortex, as well as ventral temporal cortex (Fig. 2b). The scene features yielded significant, but weaker, predictions in LOC, posterior parietal cortex (PPC), retrosplenial cortex, and ventral temporal cortex extending medially into parahippocampal cortex (Fig. 2c). Finally, the motion-energy features yielded significant performance primarily in early visual areas, including primary visual cortex (V1) and posterior VT (Fig. 2d). In general, these cortical topographies are consistent with the functional organization reported by studies using experimental manipulations to target particular specific features in isolation.

**Figure 2.**
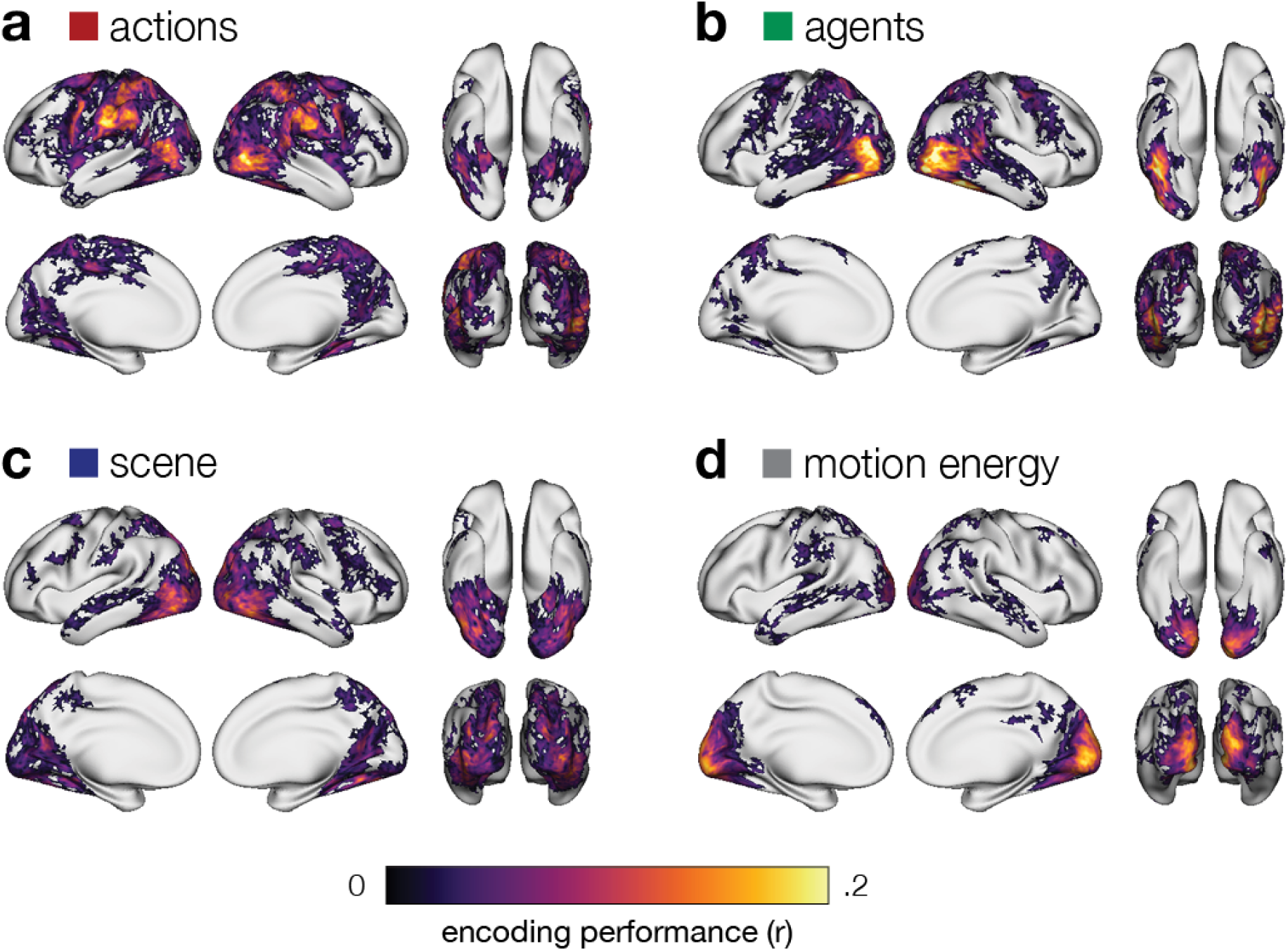
Encoding performance for each feature space. Whole-brain vertex-wise encoding performance for four feature spaces: (**a**) actions, (**b**) agents, (**c**) scene, and (**d**) motion energy. Joint encoding models were estimated combining all four feature spaces using banded ridge regression, to allow each feature space to compete for variance. Predictions were then generated from the learned weights for each feature space separately. Encoding performance is quantified as the correlation between model-predicted and actual neural activity in a left-out run in a left-out subject. Correlations are averaged across test runs and test subjects, and thresholded for statistical significance (*t*-test across subjects, FDR *p* < .05).

To simplify these results, we averaged vertex-wide encoding performance values into 10 regions of interest (ROIs). These ROIs are derived from a multimodal atlas (Glasser et al., 2016) and selected to recapitulate a number of areas identified in the literature spanning the visual processing hierarchy (Fig. 3). The action features also yielded strong predictions in LOC, but not as strong as the agent features; action features, however, were the strongest predictors in AIP and left ventral premotor cortex (vPM; Fig. 3a). The agent features yielded the strongest ROI predictions overall, localized to LOC and fusiform face cortex (FFC), with strong predictions in right-lateralized pSTS as well (Fig. 3b). Scene features performed well in occipital and temporal areas, as well as PPC (Fig. 3c). Motion energy performed best in V1, with notable prediction performance in PPC as well (Fig. 3d). Note that while action and agent features likely correlate with activity in V1, the motion-energy features (and to a lesser extent the scene features) appear to sequester much of the variance in early visual processing. On the other hand, in areas like LOC, variance in neural activity appears to be partly captured by all three semantic feature spaces.

**Figure 3.**
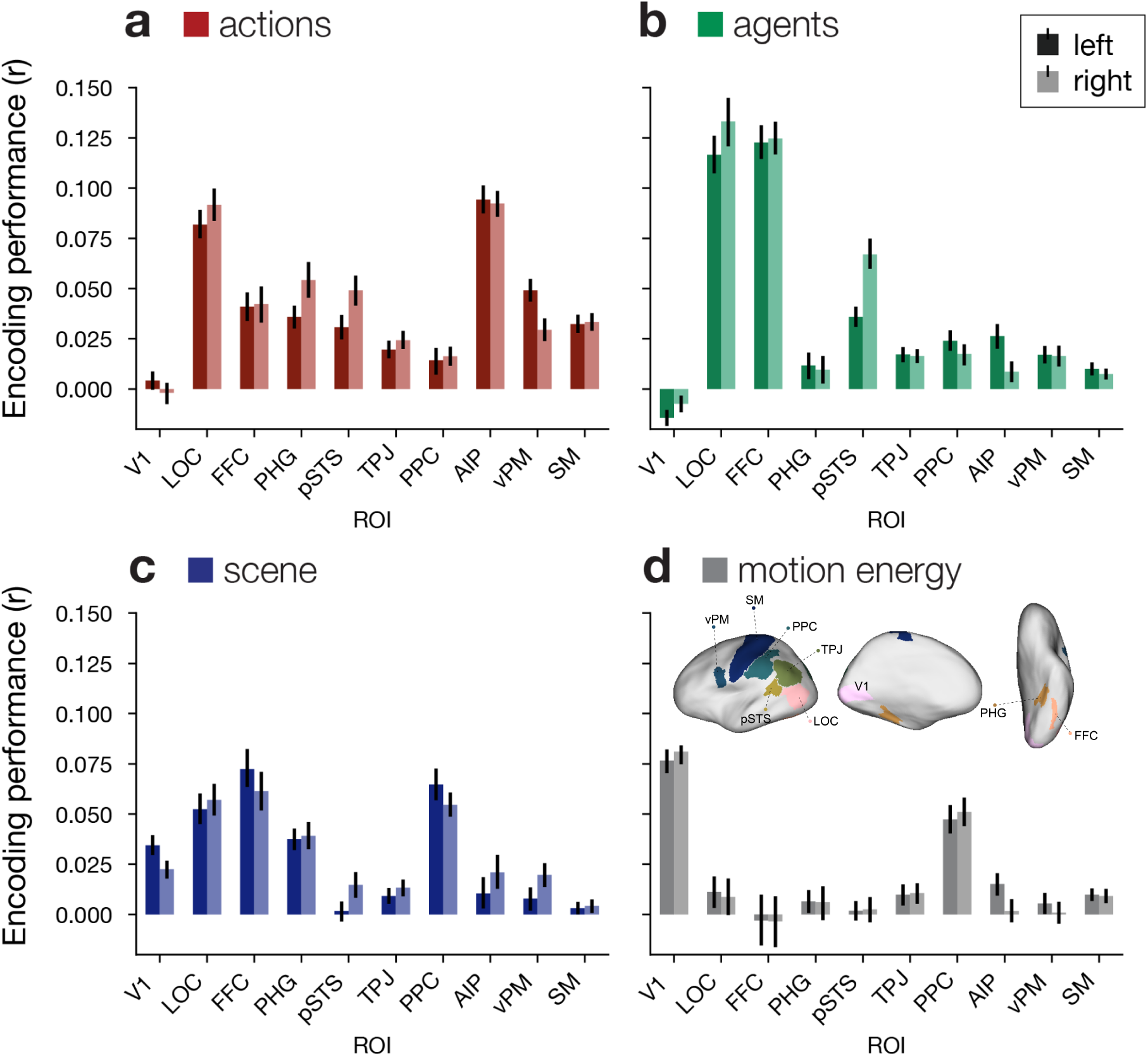
Encoding performance across regions of interest. To summarize encoding results, we averaged encoding performance values across vertices within each of 10 ROIs ranging from low-level visual areas to higher-level cortical areas. Bars indicate encoding performance averaged across left-out runs and left-out subjects for the four feature spaces: (**a**) actions, (**b**) agents, (**c**) scene, and (**d**) motion energy. Homologues in the left hemisphere (dark colors) and right hemisphere (light colors) are shown separately. Error bars indicate 95% bootstrap confidence intervals (resampling subjects).

### Comparing models for actions, agents, and scenes

We next directly compared encoding performance between each of the three types of semantic features (Fig. 4). We found that the action features significantly outperform agent and scene features throughout anterior parietal and premotor areas (paired *t*-test, FDR *p* < .05). Action features also outperformed agent features in more medial ventral temporal areas, and outperformed scene features in LOC. Agent features outperformed both action and scene features in LOC, with a strong advantage along the fusiform gyrus in ventral temporal cortex. Action and agent features performed comparably in pSTS, but both outperformed scene features. Scene features performed more strongly than actions and agents in posterior and medial visual areas and performed more strongly than agents in medial ventral temporal cortex.

**Figure 4.**
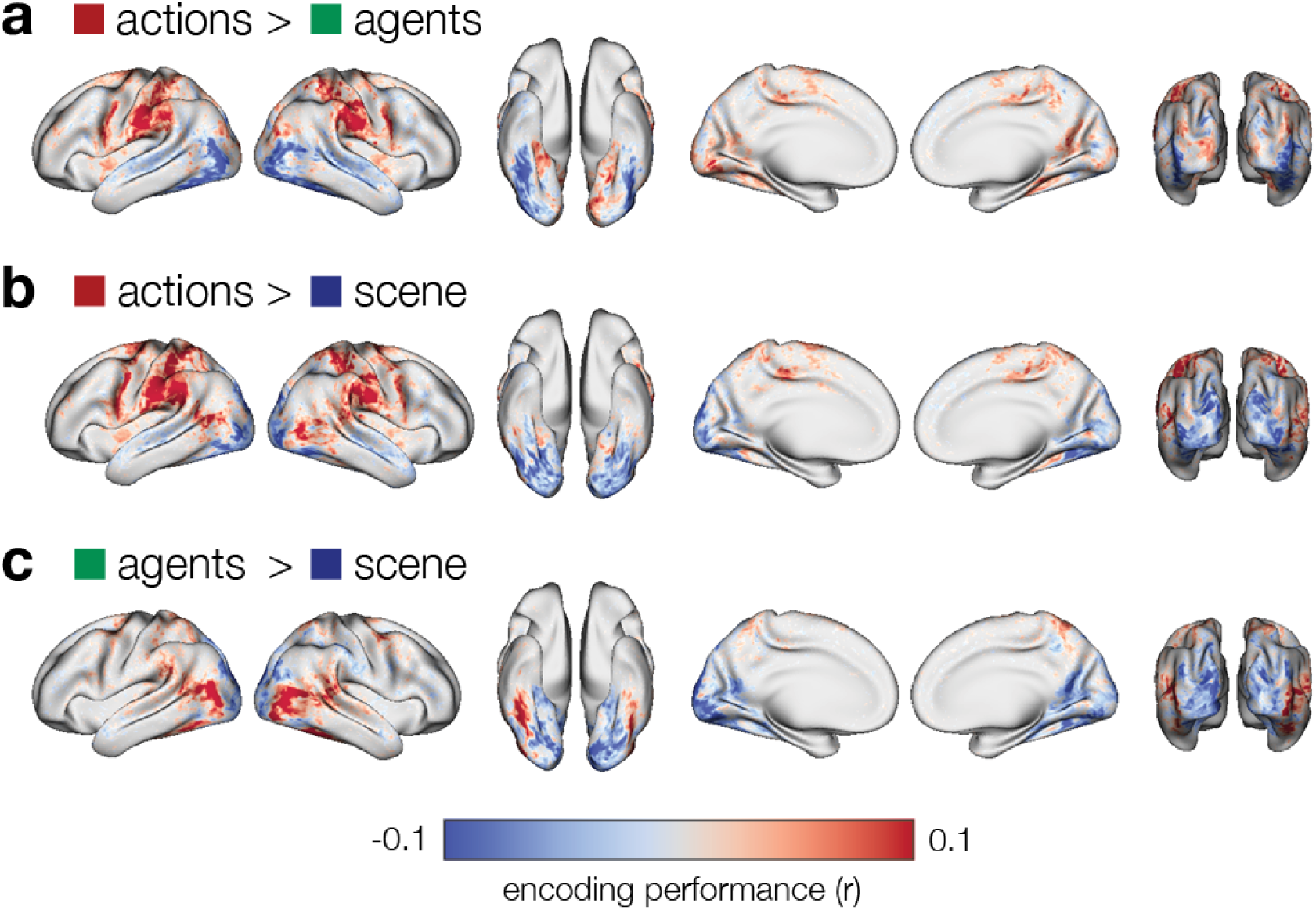
Comparing encoding performance across feature spaces. Differences in encoding performance were computed for each pair of the three feature spaces: (**a**) actions versus agents, (**b**) actions versus scene, (**c**) agents versus scene. Encoding models were estimated jointly, while predictions were generated from separate feature spaces. Differences in correlation are averaged across test runs and test subjects (paired *t*-test across subjects, FDR *p* < .05).

### Unique variance explained by each set of features

In the previous analyses, we fit a joint model with four feature bands and then generated predictions from the weights corresponding to each feature space. While this approach allows us to visualize relative contributions of different feature spaces (Lee Masson & Isik, 2021), the different feature bands may nonetheless predict overlapping portions of variance in neural activity—in a naturalistic context, different features may be highly collinear. That is, the prior analyses do not tell us how much variance in neural activity is *uniquely* predicted by a given feature space. To quantify the unique variance predicted by each feature space, we performed a variance partitioning analysis (Lescroart et al., 2015; de Heer et al., 2017). We performed a nested regression analysis in which we constructed a joint encoding model combining all four feature spaces, then fit four nested models, each excluding one feature space of interest. We quantify the unique variance explained by a feature space as the difference in encoding performance between the nested model where that feature space is excluded and the joint model where that feature space is included. This analysis effectively controls for collinearity between different sets of features; if a given feature set is collinear with other feature sets, it will not capture any unique variance in this analysis. We first quantified the unique variance explained by each of the four feature spaces and statistically evaluated these maps across subjects (Fig. 5).

**Figure 5.**
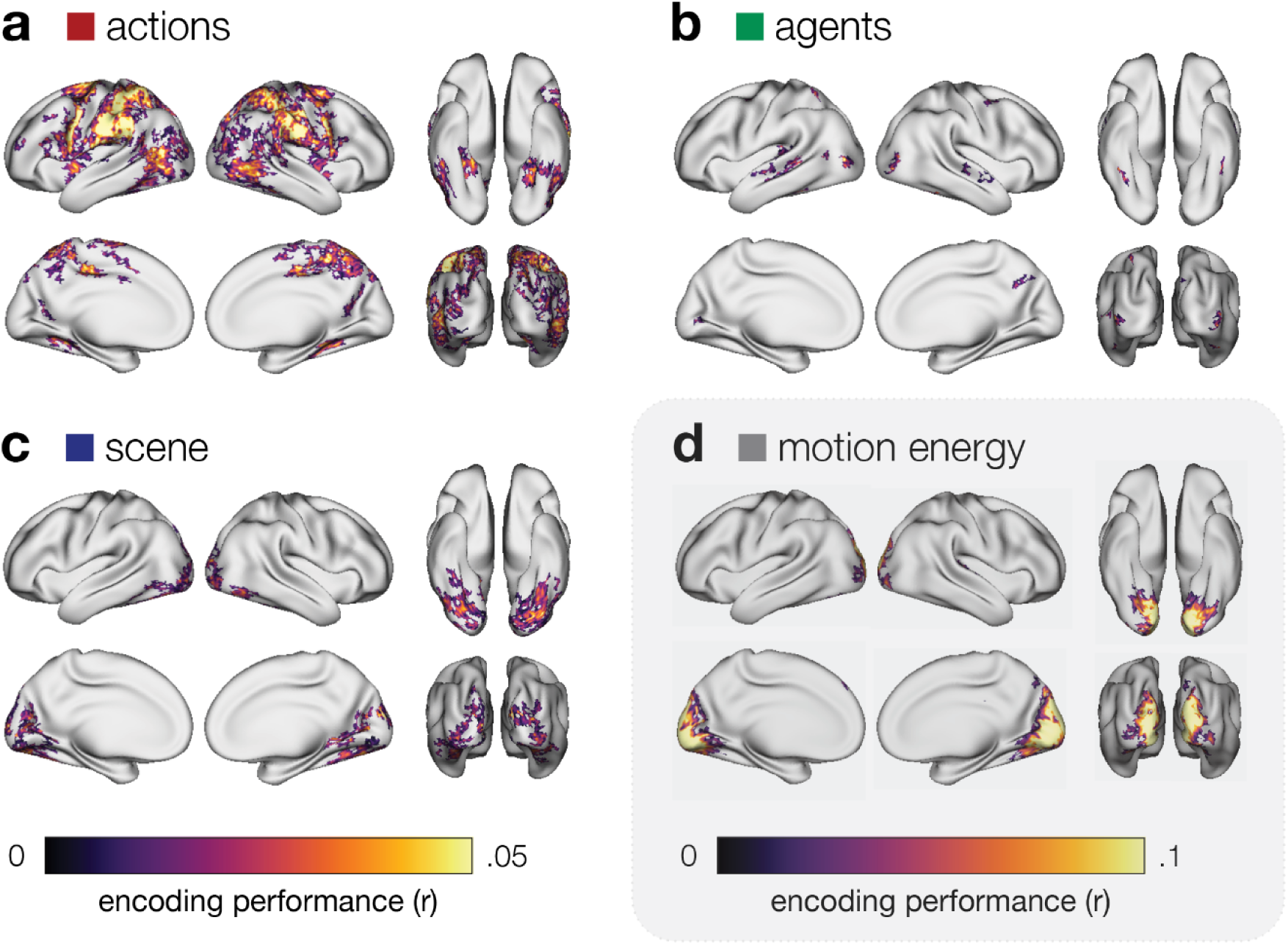
Unique variance explained by each feature space. Whole-brain maps of unique variance explained by (**a**) action, (**b**) agent, (**c**) scene, and (**d**) motion-energy features. Unique variance explained by a given feature space was estimated by computing nested regression models, then subtracting the performance of a nested model *excluding* the features of interest from a full model containing the features of interest. Vertex-wise unique variance was estimated within each run and subject, then averaged across runs and subjects, and thresholded for statistical significance (bootstrap hypothesis test resampling subjects, FDR *p* < .05).

We found that the action features uniquely predict neural activity across a large swath of cortex (Fig. 5a), including LOC, AIP, and vPM, as well anterior VT, excluding the fusiform gyrus (bootstrap hypothesis test, FDR *p* < .05). The agent features uniquely predicted a much more restricted, punctate set of vertices localized to posterior LOC, STS, and fusiform gyrus (Fig. 5b). The scene features uniquely predicted neural activity in posterior VT and medial visual areas like retrosplenial cortex (Fig. 5c). The motion-energy features captured unique variance in early visual areas (Fig. 5d).

Lastly, we sought to quantify how much of the cortical sheet was uniquely predicted by different partitions of variance across all four feature spaces. To do this, we first constructed encoding models for every combination of the four feature spaces: each individual feature space, each pair, each triplet, and the full model comprising all four feature spaces (15 combinations in total). We averaged encoding performance across subjects and runs for each combination of models. We then quantified the variance explained uniquely by each feature space, and each intersection of two or more feature spaces. This resulted in 15 values corresponding to each partition of a four-way Venn diagram. Finally, we determined which of these partitions explained the *most* variance in neural activity at each vertex (Table S1). Although we included the models containing motion-energy features in the variance partitioning analysis, we focus on the three semantic feature spaces (i.e., actions, agents, scene).

First, we found that each model uniquely predicted *some* set of vertices better than all other models (and combinations of models), suggesting that the models themselves are comparable in quality to one another. For example, agent features captured more unique variance than all other combinations in 475 vertices (Fig. 6). Critically, however, we found that action features uniquely predicted neural activity better than all other combinations of features across an order of magnitude more vertices than agent and scene features: 4,984 vertices for actions versus 475 vertices for agents and 495 vertices for scene features (Fig. 6). This indicates that a much larger proportion of human cortex is dedicated to understanding observed actions than to other commensurate features. The unique motion-energy partition, on the other hand, best explained 5,586 vertices, primarily in early visual areas, and the intersection of motion energy and semantic features best explained many vertices progressing anteriorly into extrastriate cortex. Many regions exhibited mixed selectivity for multiple semantic feature spaces. For example, 1,204 vertices, primarily in lateral and ventral occipitotemporal areas, were best explained by the intersection of all three semantic feature spaces. The intersection of actions and agents best explained 1,482 vertices, including in right pSTS. In fact, the intersection of all four feature spaces best explained the largest number of vertices overall: 15,239 vertices throughout higher-level visual areas, like VT, LOC, STS, and PPC.

**Figure 6.**
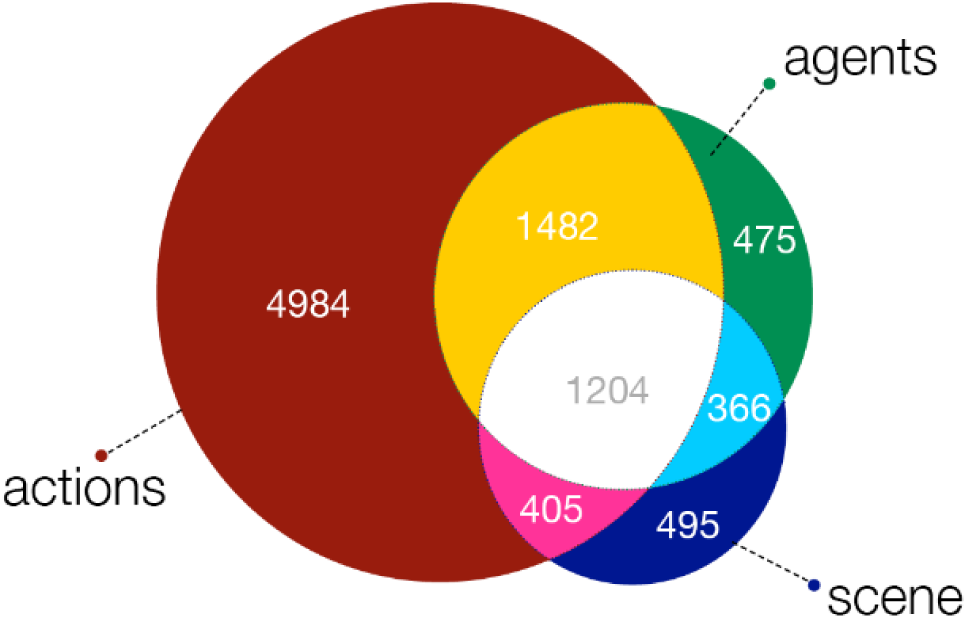
Largest partition of variance at each vertex across cortex. For each vertex, we partitioned the variance explained into components comprising the unique contributions of each feature space and the intersections of two or more feature spaces. We then assessed which of these partitions corresponded to the largest component of variance at each vertex. The Venn diagram depicts, for each partition, the number of vertices for which that partition was larger than all other partitions. For example, in 4,984 vertices, action features uniquely accounted for more variance in neural activity than any other partition. The motion-energy model was included when quantifying unique variance explained, but is excluded for visualization purposes.

## Discussion

Human cortex prioritizes certain ecologically relevant features of the world for processing. Natural vision is a holistic family of processes conspiring to understand natural scenes, their constituent agents and objects, and the behavior of those agents, all in service of our behavioral goals (Peelen & Downing, 2017; Bracci & Op de Beeck, 2023). Our understanding of the functional organization of cortex, however, stems from experiments designed to isolate particular features, typically using highly-controlled experimental stimuli (e.g., Saxe et al., 2006; Kanwisher, 2010). In the current study, we used a naturalistic movie-viewing paradigm combined with a specialized modeling framework to more fairly assess the relative contributions of different features to cortical activity. We tested three different sets of semantic features reflecting the agents, actions, and scenes depicted in the stimulus, as well as a model of low-level visual motion. We found that the actions model in particular provided strong predictions across a large portion of cortex, extending from occipitotemporal areas into parietal and frontal cortices. The action features captured unique variance in neural activity across an order of magnitude more cortical vertices than the agent or scene features. Our findings indicate that observed actions occupy a privileged role in cortical processing.

Each of the four models we tested outperformed the other models for *some* cortical areas (Figs. 2–4), suggesting that the different features are relatively well-matched; no feature set is degenerate or completely redundant with the others (see also Fig. S1). The motion-energy model was the best model in early visual areas (Nishimoto et al., 2011), but was overtaken by the semantic models in higher-level areas (Huth et al., 2012). When considering the relative contributions of different semantic features, we found that agent features were strongly encoded in ventral temporal cortex, particularly the fusiform gyrus (Kanwisher et al., 1997; McCarthy et al., 1997; Kanwisher & Yovel, 2006), and the right pSTS (Said et al., 2010; Carlin et al., 2011; Pitcher et al., 2011; Deen et al., 2015). Agent features were also a strong predictor in LOC, yielding the highest encoding performance across all models and areas in the ROI analysis (Fig. 3b). Agent features may co-vary with face (Gauthier et al., 2000; Haxby et al., 2000), body-part (Downing et al., 2001; Orlov et al., 2010; Bracci et al., 2012), and social (Wurm et al., 2017) features encoded in lateral occipital and temporal areas. Scene features performed well in ventral temporal areas, including parahippocampal cortex (Epstein & Kanwisher, 1998; Bonner et al., 2016), as well as posterior occipitoparietal cortex (Dilks et al., 2013), and retrosplenial cortex (Vann et al., 2009; Park et al., 2015). Action features performed well throughout the action observation network, including LOC (Decety & Grèzes, 1999; Kable et al., 2002, 2005; Kalénine et al., 2010; Lingnau & Downing, 2015; Wurm et al., 2017), as well as anterior parietal and premotor areas (Grafton & Hamilton, 2007; Caspers et al., 2010; Rizzolatti & Sinigaglia, 2010; Oosterhof et al., 2013). In general, our findings align with previous controlled experiments, but extend these trends of cortical functional organization into a more naturalistic setting.

For the preceding results, we evaluated each feature space after fitting all features jointly, allowing each set of features to compete for variance in neural activity (Lee Masson & Isik, 2021; Dupré la Tour et al., 2022). Although that approach allows us to assess the relative contributions of each feature space to joint encoding performance, it does not allow us to quantify how much variance in neural activity is *uniquely* predicted by each set of features. To quantify the unique variance explained by each set of features—more strictly controlling for all other features, including motion energy—we used a variance partitioning analysis (Lescroart et al., 2015; de Heer et al., 2017) (Fig. 5). We found that, while the agent and scene features still accounted for unique variance in expected regions, cortical coverage was much more focal. For example, cortical vertices uniquely captured by scene features were relatively restricted to posterior ventral temporal regions, occipitoparietal cortex, and medial visual areas like parahippocampal cortex and retrosplenial cortex. Cortical vertices uniquely captured by agent features were even sparser, with localized clusters of vertices in the bilateral fusiform gyrus, lateral occipital cortex, and STS. Action features, on the other hand, captured unique variance across a much larger portion of cortex, including VT, medial parietal cortex, and canonical action observation areas extending from LOC to AIP and VPM.

One of the principal challenges of naturalistic neuroscience stems from the inherent collinearity among stimulus features in real-world contexts. For example, agent and action features tend to co-occur in natural vision, leading to intertwined features in the regression model. As a result, many cortical areas can be predicted by both sets of features (Fig. 2a and 2b). However, under a more strict variance partitioning regime, we find that the agent features perform poorly and much of this variance is assigned to the action features instead (Fig. 5a and 5b). This observation suggests that agent features (and to a lesser extent scene features) are correlated with action features, but that neural activity predicted by agent and scene features tends to be better explained by action features. By including motion-energy features in our variance partitioning analysis, we ensure that this effect is not driven by low-level visual motion. That said, our variance partitioning analysis also revealed that a large portion of cortical vertices exhibit mixed selectivity: activity at these vertices is best explained by the intersection of different feature spaces (Table S1).

Although our encoding results recapitulate familiar maps of cortical functional organization, the relative differences in variance explained are instructive. Our findings indicate that, during natural vision, roughly 10 times as much of cortex is allocated to representing observed actions as compared to agents and scene features (Fig. 6). When viewing agents behaving in their natural environments, the “behavior” appears to be what matters most to cortex. This aligns with recent work suggesting that the dynamic features of observed actions play an outsized role in cortical activity, including in unexpected cortical areas like ventral temporal cortex (Russ & Leopold, 2015; Nastase et al., 2017; Han et al., 2024). We contend that this primacy of action representation in vision has been historically underappreciated due to the field’s reliance on highly-controlled experimental manipulations and static image stimuli (Haxby, Gobbini, et al., 2020; Leopold & Park, 2020; Nastase et al., 2020).

Our findings raise a difficult question: Why are cortical resources so heavily skewed toward understanding the actions of others? Why are these action features “magnified” across the cortical surface? While processing natural scenes and the agents occupying them is certainly important, the actions of an agent are likely to change most meaningfully over time. As an agent pursues its goals, its behavior dynamically evolves over time—these actions are laden with intentions, oftentimes social content, and sometimes behavioral ramifications for the observer. Humans are a remarkably social species and understanding the actions and intentions of others is the bedrock of social cognition.

From an ecological perspective (Gibson, 1979), all of these features of the world afford certain adaptive behaviors and may “compete” to capture cortical resources over the course of evolution and development (Zador, 2019; Hasson et al., 2020; Arcaro & Livingstone, 2021). We speculate that the actions of other agents are particularly important for our own behavior, and have therefore secured a remarkably large proportion of cortical real estate.

## Methods

### Participants

Eighteen right-handed adults (10 reported female, mean age ± SD: ∼25.4 ± 2.6 years) participated in the experiment. All participants had normal or corrected-to-normal vision, normal hearing, and reported no neurological conditions. All participants provided written, informed consent prior to participating in the study. The study was approved by the Institutional Review Board of Dartmouth College. These data have been analyzed in previous publications (Nastase et al., 2017; Van Uden et al., 2018).

### Stimuli

Participants viewed four audiovisual clips from the *Life* nature documentary narrated by David Attenborough during MRI scanning. The stimulus was divided into four ∼15-minute runs (15.3, 14, 15.4, and 16.5 minutes), totaling 63 minutes. The video stimulus was back-projected on a screen at the end of the scanner bore and viewed with a mirror attached to the head coil. The audio track was delivered using MRI-compatible fiber-optic electrodynamic headphones. Participants freely viewed the movie and were instructed to remain still. The stimulus was presented using PsychoPy (Peirce, 2008).

### Data acquisition

MRI data were acquired using a 3T Philips Intera Achieva MRI scanner with a 32-channel phased-array head coil. Functional images were acquired using single-shot gradient-echo echo-planar imaging sequence: TR/TE = 2500/35 ms, flip angle = 90°, resolution = 3 mm isotropic voxels, matrix size = 80 × 80, FOV = 240 × 240 mm, 42 axial slices with full brain coverage and no gap, anterior–posterior phase encoding, interleaved acquisition, SENSE parallel acceleration factor = 2, with fat suppression. Four runs were collected for each participant, consisting of 374, 346, 377, and 412 dynamic scans, or 935, 865, 942.5, and 1030 s, respectively. T1-weighted structural images were acquired using a high-resolution 3d turbo field echo sequence: TR/TE = 8.2/3.7 ms, flip angle = 8°, resolution = 0.9375 × 0.9375 × 1.0 mm, matrix size = 256 × 256 × 220, and FOV = 240 × 240 × 220 mm.

### Preprocessing

MRI data were preprocessed using fMRIPrep (1.0.0-rc5; Esteban et al., 2018). T1-weighted images were skull-stripped using ANTs (Avants et al., 2008). Cortical surfaces were reconstructed using *recon-all* in FreeSurfer (Dale et al., 1999) and aligned to the *fsaverage6* template (Fischl et al., 1999).

Brain tissue segmentation of cerebrospinal fluid, white matter and gray matter was performed using *fast* in FSL (Zhang et al., 2001). Functional data were slice-time corrected using *3dTshift* in AFNI (Cox & Hyde, 1997) and motion corrected using *mcflirt* in FSL (Jenkinson et al., 2002). The functional data were then co-registered to the corresponding T1 image using boundary-based registration with nine degrees of freedom implemented using *bbregister* in FreeSurfer (Jenkinson et al., 2002; Greve & Fischl, 2009). The motion correction transformations and functional-to-T1 transformation were applied in a single step using Lanczos interpolation. The functional data were then sampled to the surface by averaging across the cortical ribbon: at each vertex, the segment normal to the white-matter surface, extending to the pial surface, was sampled at 6 intervals and averaged. We next performed confound regression using *3dTproject* in AFNI (Cox, 1996). The confound model consisted of six head motion parameters, framewise displacement (Power et al., 2014), five principal components from a conservative mask of cerebrospinal fluid and white matter (aCompCor; Behzadi et al., 2007), first-and second-order polynomial trends, and a band-pass filter (0.00667–0.1 Hz)

### Hyperalignment

We used hyperalignment to align fine-grained functional topographies across individual brains prior to the vertex-wise encoding analysis (Haxby et al., 2011; Haxby, Guntupalli, et al., 2020). Specifically, we performed whole-brain surface-based hyperalignment using 20-mm-radius searchlights (Guntupalli et al., 2016). We estimated hyperalignment transformations from the three training runs in the same leave-one-run-out cross-validation procedure used for encoding model evaluation (see “Vertex-wise encoding models” below). The transformations derived from the training runs were applied to the training runs for vertex-wise encoding model estimation, then applied to the left-out test run for encoding model evaluation. In other words, the same training runs are used for hyperalignment estimation and encoding model estimation, and learned hyperalignment transformations and encoding models are tested on a previously unseen test run (Van Uden et al., 2018; Nastase, Liu, et al., 2020).

Prior to estimating the vertex-wise encoding models, data from the three training runs in *N* – 1 training subjects were transformed into the common space constructed by hyperalignment (visualized in the anatomical space of an arbitrary reference subject). Data from the *N* – 1 training subjects were averaged across subjects in common space, and the encoding model was trained on the averaged data. We then projected the left-out test subject into this common space, and evaluated the encoding model predictions against the test subject’s actual data in common space.

### Regions of interest

Regions of interest (ROIs) were constructed from a multimodal cortical parcellation (2016). We selected ten ROIs to reflect a range of areas across the cortical hierarchy commonly associated with visual object and action recognition. In the following, we report the parcel names and indices used to construct each ROI using the labeling convention of Glasser and colleagues (2016). Primary visual cortex (V1): Primary Visual Cortex (V1, 1); fusiform face complex (FFC): Fusiform Face Complex (FFC, 18); parahippocampal gyrus (PHG): ParaHippocampal Area 1 (PHA1, 126), ParaHippocampal Area 2 (PHA2, 155); ParaHippocampal Area 3 (PHA3, 127); lateral occipitotemporal complex (LOC): Middle Temporal Area (MT, 23), Middle Superior Temporal Area (MST, 2), Area TemporoParietoOccipital Junction 2 (TPOJ2, 140), Area TemporoParietoOccipital Junction 3 (TPOJ3, 141), Area V4t (V4t, 156), Area FST (FST, 157), Area Lateral Occipital 3 (LO3, 159); posterior superior temporal sulcus (pSTS): Superior Temporal Visual Area (STV, 28), Area TemporoParietoOccipital Junction 1 (TPOJ1, 139); temporoparietal junction (TPJ): Area PFm Complex (PFm, 149), Area PGi (PGi, 150), Area PGs (PGs, 151); posterior parietal cortex (PPC): Area V3A (V3A, 13), Area V3B (V3B, 19), Seventh Visual Area (V7, 16), IntraParietal Sulcus Area 1 (IPS1, 17); anterior intraparietal cortex (AIP): Area PFt (PFt, 116), Anterior IntraParietal Area (AIP, 117), Area PF opercular (PFop, 147), Area PF Complex (PF, 148); somatomotor cortex (SM): Primary Motor Cortex (4, 8), Primary Sensory Cortex (3b, 9), Area 1 (1, 51), Area 2 (2, 52), Area 3a (3a, 53); ventral premotor cortex (vPM): Rostral Area 6 (6r, 78), Area IFJp (IFJp, 80).

### Stimulus annotation

In order to quantify the semantic content of the movie stimulus, we first assigned word labels to each moment of the film (Huth et al., 2012; Van Uden et al., 2018). A human annotator assigned words describing the visual content of the stimulus in roughly 2-TR segments averaging 5.17±3.85 seconds (run 1 = 5.83; run 2 = 4.61; run 3 = 5.59; run 4 = 4.67; certain segments—e.g., the introductory sequence—were longer than others). The audio track and voice-over narrative were not presented during this annotation task. The words in each segment were then split into three different sets: words describing the *actions* taken by animals in the film (e.g., “swimming”), words describing the *agents* themselves (e.g., “dolphin”), and words describing the *scene* or background (e.g., “ocean”). Words from each segment were then duplicated to effectively upsample the roughly 5-second annotation windows to the 2.5-second TR of the fMRI acquisition.

### Semantic feature spaces

To quantify the semantic content of the visual stimulus, we extracted 300-dimensional semantic vectors from GloVe (Pennington et al., 2014). GloVe embeddings have been widely used to capture semantic content encoded in neural activity (e.g., Pereira et al., 2018; Anderson et al., 2021; Fernandino et al., 2022; Goldstein et al., 2022; Grand et al., 2022; Kumar et al., 2024). GloVe is an unsupervised learning algorithm that represents words as vectors in a high-dimensional embedding space based on their co-occurrence statistics in large corpora of text. Semantic structure is encoded in the geometric relationships between word embeddings in this high-dimensional vector space: certain semantic relationships are encoded along particular directions and semantically similar words are nearer to each other along certain dimensions (Landauer & Dumais, 1997; Mikolov et al., 2013).

Each word receives the same embedding regardless of its local context or usage in our annotation. For each set of annotations (actions, agents, and scene words), we compiled the embeddings into TRs and averaged embeddings for multiple words occurring within a single TR. This resulted in one embedding for actions, one embedding for agents, and one embedding for scene words per TR. Any TRs containing no words were assigned a 300-dimensional vector of zeros.

### Motion-energy model

Prior work suggests that visual motion has a large impact on cortical activity in response to dynamic, naturalistic visual stimuli like movies (Nishimoto et al., 2011; Huth et al., 2012). To account for low-level visual motion, we submitted our movie stimuli to a motion-energy model (Adelson & Bergen, 1985; Watson & Ahumada, 1985) implemented in *pymoten* (Nishimoto & Gallant, 2011; A. Nunez-Elizalde et al., 2021). Video frames were first converted to luminance values and downsampled to 192 × 108 pixels, yielding a 2,530-parameter pyramid of motion energy filters tiling the stimulus (according to the default parameters in *pymoten*). The resulting motion-energy features were log-transformed and downsampled from the original stimulus frame rate to match the TR of the fMRI data.

### Vertex-wise encoding models

To quantify which features of the movie stimulus are encoded in neural activity—e.g., low-level motion features or higher-level semantic features—we constructed vertex-wise encoding models (Naselaris et al., 2011; Huth et al., 2012). Vertex-wise encoding models estimate a linear mapping from a representation of the stimulus in a particular to feature space onto the time series of neural activity. To more equitably compare different feature spaces, we use banded ridge regression to fit all feature spaces simultaneously in a joint model (A. O. Nunez-Elizalde et al., 2019; Dupré la Tour et al., 2022).

We quantify how well these features map onto vertex-wise activity by (1) evaluating the predictions from each feature space separately and (2) evaluating the unique variance explained by each feature space (and intersection of feature spaces). In the following, we describe the vertex-wise encoding analysis in greater detail.

#### Preparation of model features

To prepare the model features for vertex-wise encoding analysis, we first split the sequence of embeddings for each feature space—actions, agents, scene, and motion energy— according to the four scanner runs, and reserved three runs for training and one run as the test set (following four-fold leave-one-run-out cross-validation). To match the dimensionality of each feature space, we applied PCA to the matrix of semantic embeddings separately for each of the four: (1) we z-scored the features of the three concatenated train runs and applied that z-transformation from the training runs to the test run; (2) we then applied PCA to a given feature space to reduce the original dimensionality (300 for GloVe embeddings, 2,530 for motion energy) to 40 components, capturing over 95% of the variance in a feature space (Fig. S1); and (3) we projected the features of the test run onto the principal components estimated from the training runs to avoid any leakage between training and test set.

To account for variable hemodynamic lags, we implemented a finite impulse response (FIR) model, duplicating and concatenating each feature space with lags of 1 TR (2.5 s), 2 TRs (5 s), 3 TRs (7.5 s), and 4 TRs (10 s). The regression model then estimates a linear combination of features across lags to best predict activity at a given vertex; this approach allows for us to capture the optimal combination of lags across features, as different features may have different temporal dynamics.

#### Preparation of fMRI data

The fMRI time series were prepared for vertex-wise encoding analysis by (1) organizing the four scanner runs into training and test sets according to a four-fold leave-one-run-out cross-validation scheme; and (2) organizing the subject data into training and test sets according to a leave-one-subject-out cross-validation scheme. The fMRI data for each subject and each run were functionally aligned into a shared space using searchlight hyperalignment, where the hyperalignment transformations were estimated from (and applied to) three training runs, and this hyperalignment transformation was then applied to the test set, following the same leave-one-run-out cross-validation scheme as used in the vertex-wise encoding analysis (see the “Hyperalignment” section above). For the training fMRI data from *N* – 1 subjects, we first (1) z-scored each vertex time series across time within each run; then (2) averaged the hyperaligned time series across subjects; then (3) z-scored the average time series within each run prior to concatenating the training runs. These hyperaligned and averaged time series derived from three training runs and *N* – 1 subjects served as the training data for the vertex-wise encoding analysis at a given cross-validation fold The test data comprised fMRI responses for the left-out scanner run in a left-out subject; the test subject’s data were z-scored across time at each vertex.

##### Estimating and evaluating vertex-wise encoding models

We used banded ridge regression to estimate a weight matrix that linearly maps from the combined features onto the fMRI activity at each vertex (A. O. Nunez-Elizalde et al., 2019; Dupré la Tour et al., 2022). Ridge regression mitigates overfitting by imposing a penalty on the magnitude of the weights. Banded ridge regression allows us to fit multiple feature spaces in the same encoding model, where each feature space receives its own penalty term. We implemented a four-fold leave-one-run-out and leave-one-subject-out outer cross validation scheme to prevent data leakage and robustly evaluate the out-of-sample prediction performance of our models. Across the four concatenated feature bands—actions, agents, scenes, and motion energy—we estimate a single weight vector mapping from the features onto the fMRI time series at a given vertex. To optimize the four penalty terms corresponding to the four feature spaces, we used a random search (1000 iterations) across combinations of penalty terms with nested three-fold leave-one-run-out cross-validation within the training set of the outer cross-validation loop. For a single outer cross-validation fold, this resulted in one weight matrix across all vertices derived from three training runs and N – 1 training subjects. We then applied this weight matrix to the model features from the test run, yielding model-predicted fMRI time series for the test run. To quantify the model performance, we computed the Pearson correlation coefficient between the model-predicted time series and the actual fMRI time series for the left-out test subject. In this way, the generalization performance of our models was always evaluated on different stimulus content (i.e., left-out scanner runs) and different individual brains (i.e., left-out subjects) (Van Uden et al., 2018; Nastase, Liu, et al., 2020).

To evaluate the relative contributions of each feature space—actions, agents, scene, and motion energy—we generated model-based predictions using the regression weights for each feature band separately (effectively zeroing out weights for other bands) (Lee Masson & Isik, 2021). We then computed the correlation between predicted time series and actual fMRI time series in the left-out run and left-out test subject for each feature space. This results in four separate encoding performance maps, one for each feature space.

### Variance partitioning

We performed two separate variance partitioning analysis: (1) we computed the unique variance explained by each of the four models and assessed the statistical significance of these unique variance values (Fig. 5); (2) we computed all 15 partitions comprising the unique contributions and intersections between two or more sets of features, then determined which of these 15 partitions was largest for each vertex (Fig. 6). In the first analysis, we fit a full model comprising all four sets of features, then fit nested models excluding the feature space of interest. For a given feature space of interest, we quantified the unique variance explained by that feature space as *r*_unique_ = *r*_full_ *– r*_nested_, where *r*_full_ corresponds to the encoding performance for the full model with all four sets of features, and *r*_nested_ corresponds to the encoding performance for the nested model excluding the model of interest. We computed these vertex-wise *r*_unique_ values for each subject, then assessed statistical significance across subjects, resulting in a whole-brain map of statistically significant *r*_unique_ values for each of the four feature spaces (Fig. 5).

In the second analysis, we fully partitioned the variance explained by each combination of feature spaces using commonality analysis (Mood, 1971; Seibold & McPhee, 1979; Lescroart et al., 2015; de Heer et al., 2017). We ran encoding models for all possible combinations of feature spaces, resulting in 15 models (e.g., single features, pairs, triplets, and all four features: _4_C_1_ + _4_C_2_ + _4_C_3_ + _4_C_4_). We evaluated these models in the same way as previous models, by computing the correlation between model-predicted and actual vertex-wise time series (Lee Masson & Isik, 2021) for left-out runs in left-out subjects. To simplify the commonality analysis, for each of the 15 models, we first averaged correlations (with z-transformation) across both runs and subjects, to obtain a single encoding performance score for each model.

We used set theory to arithmetically determine the variance assigned uniquely to each of the four feature spaces as well as each intersection of two, three, or all four features for every vertex. This allowed us to quantify the variance uniquely explained by one feature space by isolating the variance specific to that feature space while excluding all other feature spaces and their intersections. In keeping with prior work (Lee Masson & Isik, 2021), we quantified unique variance explained in terms of the out-of-sample correlation between predicted and actual time series.

Finally, we identified which partition—i.e., unique feature space or intersection of feature spaces—explained the most variance in vertex-wise activity. This allowed us to quantify, for example, the number of vertices best explained uniquely by action features across cortex, or the number of vertices best explained by the intersection of action and motion-energy features. We computed these vertex counts within a mask of vertices with significant model performance for the joint model combining all four feature spaces (39,031 significant vertices in the mask in total across both hemispheres; bootstrap hypothesis test, FDR *p* < .05, Fig. S2). Motion energy features were included in the variance partitioning analysis, although we focus primarily on the three semantic feature sets. Note that vertices identified in the preceding analyses as having significant unique variance explained by a given feature space (Fig. 5) may be better explained by a different combination of features in this second “winner-take-all” analysis (Fig. 6).

### Statistical evaluation

To determine whether the mean encoding performance (correlation between predicted and actual response time series) across subjects was statistically greater than zero, we performed a one-sample *t*-test. To compare encoding performance between models, we submitted the subject-level correlation values for both models to a paired *t*-test. Encoding performance values were averaged across the four left-out runs (four different segments of the film) within each subject prior to statistical evaluation.

Given that the unique variance estimates are biased toward positive values due to the nested structure of the models, we used a bootstrap hypothesis test to more conservatively assess significance (Hall & Wilson, 1991). To correct for multiple tests across vertices, we controlled the false discovery rate (FDR; Benjamini & Hochberg, 1995) at .05 across both hemispheres (excluding vertices in the medial wall). To simplify visualization of whole-brain model performance, we omitted any clusters of significant vertices containing fewer than 100 contiguous surface vertices (Figs. 2, 4); when visualizing unique variance explained (Fig. 5), we omitted clusters with fewer than 50 significant contiguous surface vertices.

## Acknowledgements

We would like to thank Courtney Rogers, Terry Sackett, and the Dartmouth Brain Imaging Center for help with data collection, as well as Zaid Zada and Vassiki Chauhan for help with modeling and visualization.

## Supplementary Information

**Figure S1.**
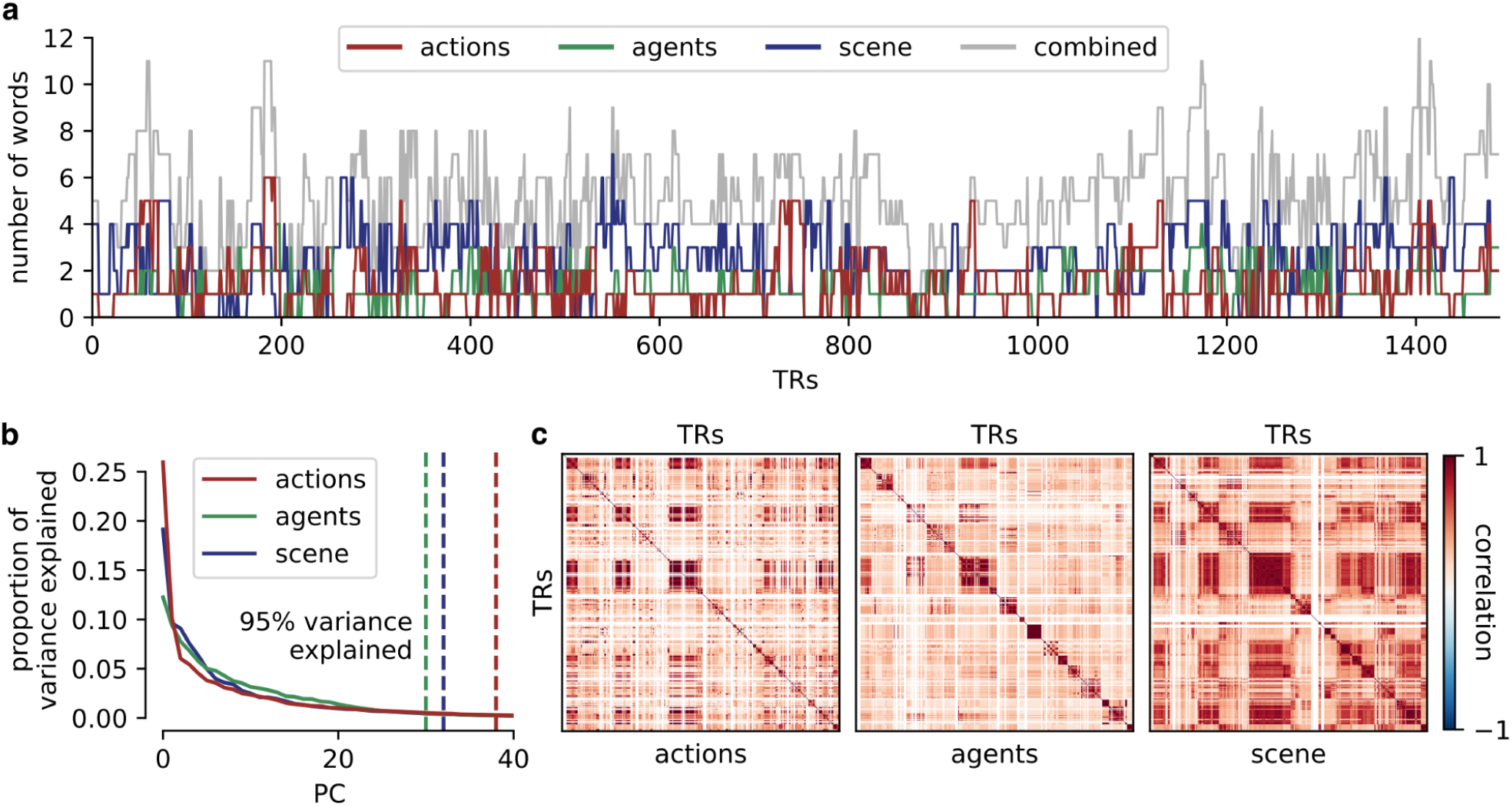
Similar frequency and dimensionality of different feature spaces. (**a**) Annotation word count per TR across runs for all three semantic feature spaces (actions, agents, and scene words). All three semantic feature spaces occurred with similar frequency across all four runs. (**b**) To estimate the dimensionality of each semantic feature space, we applied PCA to the 300-dimensional GloVe vectors assigned to each TR concatenated across all four runs. Scree plot depicts the proportion of variance explained by an increasing number of PCs. Vertical dotted lines indicate the number of PCs accounting for 95% of variance in the time series of embeddings (actions: 38 PCs, agents: 30 PCs, scene: 32 PCs). Overall, the three semantic feature spaces had similar dimensionality. (**c**) We visualized time-point (TR-by-TR) similarity matrices for each semantic feature space by computing the correlation between embeddings for a given feature space among all pairs of TRs.

**Figure S2.**
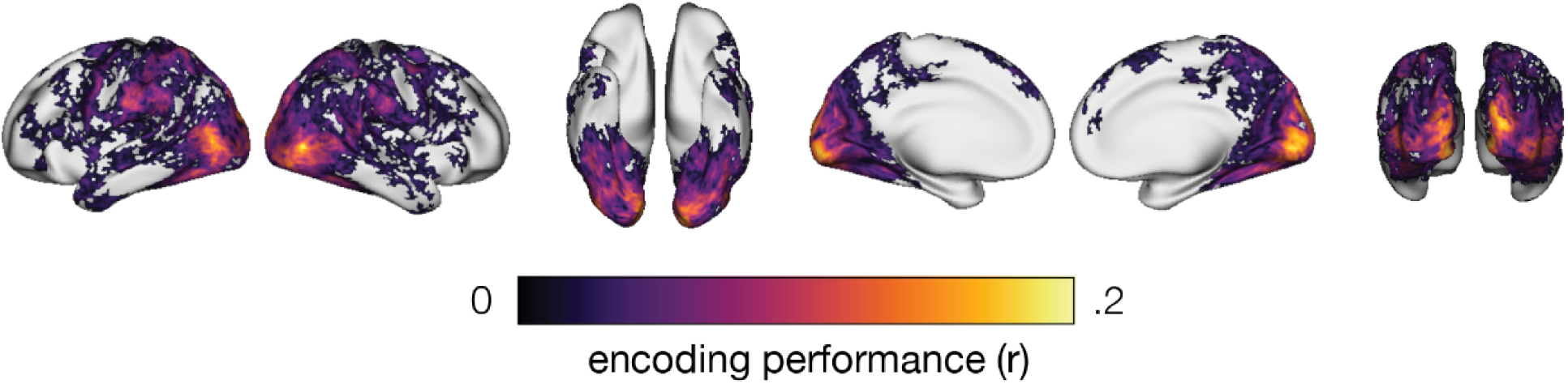
Encoding performance for the full model. For each vertex, we estimated a joint encoding model combining the four feature spaces: actions, agents, scene, and motion energy features. Here we report the encoding performance for the full model (compare to the encoding performance for individual feature bands in Fig. 2). This map was also used as a mask to identify vertices with significant encoding performance for the variance partitioning analysis (Fig. 5d). Encoding performance is quantified as the correlation between model-predicted and actual neural activity in a left-out run in a left-out subject. Correlations are averaged across test runs and test subjects, and thresholded for statistical significance (*t*-test across subjects, FDR *p* < .05).

**Table S1.** Number of cortical vertices best explained by each partition of variance. For each cortical vertex, we partitioned the variance explained by the joint model into 15 partitions corresponding to the variance explained uniquely by each model and by each intersection of two or more models (left column). For each vertex, we then computed which of the 15 partitions explained the most variance. In the table, we report the number of cortical vertices best explained by each of the 15 partitions.

| Variance partition | Left hemisphere | Right hemisphere | Total |
| --- | --- | --- | --- |
| unique action | 2872 | 2112 | 4984 |
| unique agent | 253 | 222 | 475 |
| unique scene | 239 | 256 | 495 |
| unique motion | 2585 | 3001 | 5586 |
| action ∩ agent | 726 | 756 | 1482 |
| action ∩ scene | 153 | 252 | 405 |
| action ∩ motion | 715 | 801 | 1516 |
| agent ∩ scene | 187 | 179 | 366 |
| agent ∩ motion | 3 | 4 | 7 |
| scene ∩ motion | 454 | 551 | 1005 |
| action ∩ agent ∩ scene | 615 | 589 | 1204 |
| action ∩ agent ∩ motion | 1794 | 1763 | 3557 |
| action ∩ scene ∩ motion | 1258 | 1395 | 2653 |
| agent ∩ scene ∩ motion | 22 | 35 | 57 |
| action ∩ agent ∩ scene ∩ motion | 7318 | 7921 | 15239 |

## Notes

### Competing Interest Statement

The authors have declared no competing interest.

